# Differential functions of multiple Wnts and receptors in cell polarity regulation in *C. elegans*

**DOI:** 10.1101/2025.11.25.690575

**Authors:** Hitoshi Sawa, Masayo Asakawa, Takefumi Negishi

## Abstract

Metazoan species possess multiple Wnt ligands and receptor genes that regulate diverse developmental processes. Because these genes often act redundantly, analysis of single-gene mutants does not necessarily reveal the full roles of Wnt signaling. In *C. elegans*, three Wnt genes (*cwn-1*, *egl-20*, and *cwn-2*) and three receptor genes (*lin-17*/Fzd, *mom-5*/Fzd, and *cam-1*/Ror) redundantly regulate the polarity of asymmetrically dividing seam cells. Here, we comprehensively analyzed genetic interactions among these Wnt and receptor genes. In *mom-5* mutant backgrounds, additional mutations in Wnt genes disrupted cell polarization. In contrast, in *cam-1* mutant backgrounds, Wnt mutations frequently caused abnormal polarity orientation. These findings indicate that MOM-5 and CAM-1 play distinct roles in establishing cell polarization and determining its orientation, respectively. *lin-17* mutations suppressed polarity reversal in multiple Wnt compound mutants, suggesting that LIN-17 may function as a molecular switch for polarity orientation. Although all three Wnt genes regulate polarity orientation in a gradient-independent manner in the absence of receptor mutations, in *lin-17* mutant backgrounds, reversing the expression gradients of *cwn-1* and *egl-20*, but not *cwn-2*, enhanced polarity reversal. This suggests that *cwn-1* and *egl-20* act not only permissively but also instructively to regulate polarity orientation. Together, our results reveal distinct and cooperative functions of multiple Wnt ligands and receptors that ensure robust control of cell polarity.

## Introduction

Wnt proteins play essential roles in animal development and physiology (Nusse and Clevers 2017; Rim et al. 2022). Vertebrates possess a large number of Wnt genes (e.g., 19 in humans) and receptor genes (e.g., 10 Frizzled receptors in humans), many of which may function uniquely or redundantly (Logan and Nusse 2004; Wang et al. 2016; Zheng and Sheng 2024) (The Wnt Homepage https://wnt.stanford.edu/). In mice, although single mutants of most Wnt genes exhibit developmental abnormalities, the redundancy among Wnt genes and the practical difficulty of generating compound mutants have made it nearly impossible to obtain a complete picture of Wnt functions.

Proper animal development requires coordination between cell fate specification and cell polarity. Gradients of Wnt proteins are well known to act as positional cues that organize cell differentiation through concentration-dependent effects on cell fates (Rogers and Schier 2011). Wnts also regulate cell polarity, primarily through the planar cell polarity (PCP) signaling pathway, which involves the asymmetric localization of signaling molecules such as the Wnt receptor Frizzled (Fzd) and the cytoplasmic protein Dishevelled (DVL)(Yang and Mlodzik 2015). Whether Wnt gradients also serve as positional cues for cell polarity, however, remains controversial. In some contexts, Wnts have been shown to play permissive (gradient-independent) roles in regulating polarity (Heisenberg et al. 2000; Čapek et al. 2019). Although instructive (gradient-dependent) Wnt functions have been supported by experiments in which uniform Wnt expression perturbs cell polarity (Gao et al. 2011; Minegishi et al. 2017), such effects may result from overexpression rather than true disruption of Wnt gradients. If Wnts indeed function instructively, it remains unclear how a single cell can interpret shallow and potentially fluctuating Wnt gradients.

In *C. elegans*, most mitotic cells are polarized along the anterior‒posterior (A‒P) axis and divide asymmetrically. This polarity is regulated by the Wnt/β-catenin asymmetry pathway, in which Wnt signaling components are asymmetrically localized̶β-catenin and APC to the anterior, and Frizzled (Fzd) and Dishevelled (DVL) to the posterior side of each cell (Mizumoto and Sawa 2007b; Sawa 2012). Although the involvement of the core planar cell polarity (PCP) components Van Gogh (Vangl) and Prickle has not been demonstrated, the mechanism resembles PCP in that DVL and Fzd exhibit asymmetric localization. It has been shown that Wnts instructively control the polarity of cells positioned near Wnt-expressing cells (Goldstein et al. 2006). However, it remains unknown whether Wnt gradients instruct the polarity of cells located farther from the Wnt sources.

We previously showed that the polarity of seam cells (V1‒V6), located along the lateral side of *C. elegans*, is redundantly regulated by three Wnt proteins: CWN-1 and EGL-20, expressed posteriorly near the anus and in posterior body-wall muscle, respectively, and CWN-2, expressed anteriorly in the pharynx (Yamamoto et al. 2011). Each single and double mutant exhibited only minor polarity defects, except for the V5 cell, which was affected by *egl-20* single mutations. In contrast, *cwn-1 egl-20 cwn-2* triple mutants (hereafter referred to as *3xWnt*) displayed randomized polarity orientation. Three Wnt receptors̶LIN-17/Fzd, MOM-5/Fzd, and CAM-1/Ror̶also appear to function redundantly. When Wnt gradients were reversed by expressing CWN-1 anteriorly or CWN-2 posteriorly, polarity defects in *3xWnt* mutants were suppressed, suggesting that Wnt functions in this context are permissive. Here we analyzed compound mutants carrying combinations of Wnt and receptor mutations to uncover the genetic interactions between individual Wnts and receptors. We found that each receptor exhibited distinct features depending on specific Wnts in the regulation of cell polarization̶defined here as the establishment of cell polarity irrespective of its orientation̶and polarity orientation. Moreover, reversing the gradients of the posteriorly expressed Wnts (CWN-1 and EGL-20) in the *lin-17*/Fzd mutant background enhanced polarity reversal in *3xWnt* mutants, suggesting that these Wnts act instructively as well as permissively. In contrast, reversing the CWN-2 gradient rescued the polarity defects even in *lin-17 3xWnt* mutants, demonstrating distinct roles of the three Wnts in polarity regulation.

## Materials and Methods

### Strains and analyses of polarity

The following alleles were used: *cwn-1(ok546)* (deletion); *cwn-2(ok895)* (deletion) (Zinovyeva and Forrester 2005); *egl-20(n585)* (missense, but behaves like null) (Maloof et al. 1999); *lin-17(n3091)* (nonsense) (Sawa et al. 1996); *mom-5(os171)* (see below); *cam-1(gm122)* (nonsense); *cam-1(ks52)* (deletion) (Forrester et al. 1999); *dsh-1(ok1445)* (deletion) (Klassen and Shen 2007); *dsh-2(or302)* (deletion) (Hawkins et al. 2005); *mig-5(tm2639)* (deletion); *mig-5(cp385)* (mNG::AID::*mig-5*) (Heppert et al. 2018); *osTi8* and *osTi9* (see below); *ieSi57*(*eft-3p*::TIR1::mRuby) (Zhang et al. 2015); *osIs47* (a multicopy integrant of scm promoter::LIN-17::GFP and pUnc-76); *osIs112* (a multicopy integrant of *ceh-22* promoter::EGL-20::mNG and pUnc-76); *osEx395* (an extrachromosomal array containing *ceh-22* promoter::CWN-1::Venus) (Yamamoto et al. 2011); *osIs213* (a multicopy integrant of *egl-20* promoter::mNG::CWN-2 in which mNG was inserted immediately after the signal sequence of CWN-2). The strains used in the study are listed in Table S1. The genotypes of compound strains were confirmed either by PCR (*cwn-1*, *cwn-2*, *mom-5(os171)*), sequencing or complementation tests (*egl-20, cam-1*), or by their phenotypes (Psa and Bivulva for *lin-17*). Some of the strains were maintained as heterozygotes over balancer chromosomes marked by the GFP or mKate2 expression. Non-fluorescent homozygotes were analyzed for their phenotype. The animals were cultured at 22.5°C. All the strain used in this study contains *elt-3p*::GFP (*vpIs1*) (Koh and Rothman 2001). The polarity of seam cell divisions was analyzed at the late L1 stage by confocal microscopes (Zeiss LSM510 or LSM700) using the *elt-3p*::GFP expressed in hyp7. Statistical analysis was performed with the Fisher exact test.

### AID degron experiments

*mom-5(os171)* (*mom-5*::mAID::mClover) was generated by CRISPR/Cas9 mediated genome editing according to (Dickinson et al. 2013). The plasmid containing Cas9 and sgRNA against the C-terminus of *mom-5* was same as used for *mom-5(cp367)* (mNG insertion) (Heppert et al. 2018). The repair template plasmid is derived from pDD287 and contains homology arms used for *mom-5(cp367)* (Heppert et al. 2018) and the mAID-mClover sequence from pMK290 (Natsume et al. 2016; Negishi et al. 2019 Dec 10). *osTi8* and *osTi9* contain *eft-3p*::TIR1::mRuby derived from pLZ31 (Zhang et al. 2015) and integrated near the right end of chromosome II and the center of chromosome I, respectively, using the miniMos system (Frøkjær-Jensen et al. 2014). For the auxin treatments, late-stage embryos containing *mom-5(os171) or mig-5(cp385)* and *eft-3p*::TIR1::mRuby (either *osTi8, osTi9 or ieSi57)* were placed on plates containing 250μM indole-3-acetic acid (IAA) dissolved in DMSO.

### Genome editing of *cwn-2*

We were unable to detect fluorescence in animals carrying a knock-in of mNeonGreen (mNG) at the C terminus of the endogenous *cwn-2* locus *(os172)* (data not shown). In addition, *lin-17*(*n3091 null*) *cwn-2(os172)* animals exhibited the missing-gonad-arm phenotype (42% missing anterior and 8% missing posterior arms), comparable to that observed in *lin-17(n3091) cwn-2(ok895 null)* mutants (45% and 6%, respectively) (So et al. 2024), indicating that CWN-2 with a C-terminal fusion loses its function. In contrast, fluorescence was detected in animals carrying an N-terminal mNG knock-in immediately downstream of the signal sequence (*os187)*. *lin-17(n3091) cwn-2(os187)* animals showed only weak gonadal defects (4% missing anterior and 4% missing posterior arms), as observed in *lin-17(n3091)* single mutants (6% and 1%, respectively), indicating that CWN-2 with an N-terminal fusion retains its function. To insert mNG after the signal sequence of *cwn-2*, homologous repair template was constructed by inserting PCR products of homology arms and mNG sequence into plasmids containing a self-excising selection cassette pDD287 (Addgene #70685) (Dickinson et al., 2015) using NEBuilder (NEB). Primers for the homology arms upstream and downstream of the *cwn-2* signal sequence are as follows. Left arm: AACGACGGCCAGTCGCTTGCCTGTTTCAAAGCAGAAA and TCCCTTGGAGACCATTGATTGAACATTTAATAAATTA.

Right arm: AGCGAGGAAGACTTGTTATTAGATGCTTCTTGGTGGT and CTATGACCATGTTATGTGTGTTGAACAATTCCATCGA. A sgRNA plasmid was constructed by inserting guide RNA sequence (TGATTCCACGGAGAAGTTGTTGG) into PU6::unc-119_sgRNA (Addgene #46169) using NEBuilder. Characterization of *os187* fluorescence will be reported elsewhere.

## Results

### Redundant functions of three Wnt receptors

We determined seam cell polarity by assessing the asymmetric fates of their daughter cells using *elt-3p*::GFP, which is expressed in hyp7 but not in seam cells (Gilleard et al. 1999; Koh and Rothman 2001) (Figure 1A, B). Approximately one hour after division of seam cells (V1‒V6) in wild-type L1 larvae, the anterior daughters fused with hyp7 and immediately began fluorescing by incorporating GFP from the hyp7 cytoplasm. Thus, daughter cell fates could be unambiguously identified, allowing us to deduce the division polarity type̶normal, reversed, or lost (Figure 1B). In Figure 1C and subsequent figures, the proportions of these polarity types in individual seam cells were mathematically converted to RGB colors as described in the figure legend.

**Figure 1.**
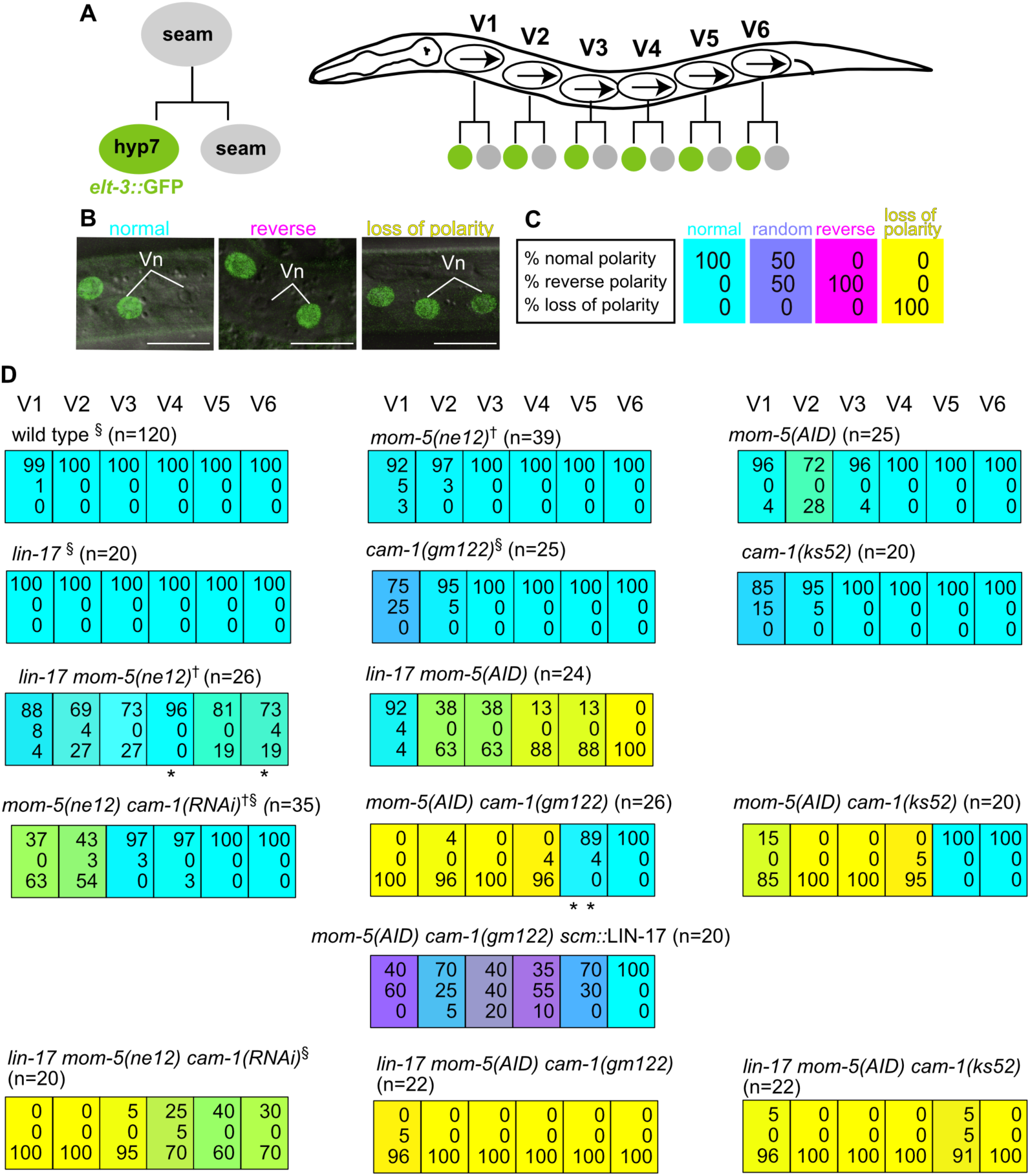
Redundant functions of three Wnt receptors. (A) Diagram of seam cell divisions. Left: The anterior daughter of a seam cell fuses with hyp7 and becomes *elt-3p*::GFP positive. Right: Seam cells (V1‒V6) along the lateral sides of the animal are polarized in the same orientation (arrows) and divide asymmetrically. (B) Examples of seam cell (Vn) division polarity̶normal, reversed, or loss of polarity-based on *elt-3p*::GFP expression in the daughter cells. Merged differential interference contrast (DIC) and fluorescence images are shown. (C) Illustration of representative polarity data for seam cell divisions. Each colored box represents one seam cell. The top, middle, and bottom numbers indicate the percentages of cells with normal, reversed, and loss of polarity, respectively. RGB color components were assigned to each phenotype (normal = red, reversed = green, loss of polarity = blue), and box colors represent the combined intensities. RGB intensity was calculated as 255 − (observed phenotype × 2.55), where 255 is the maximum component value and 2.55 converts percentages to the RGB scale. The resulting standard colors are cyan (100 percent normal), lavender (random polarity), magenta (100 percent reversed), and yellow (100 percent loss of polarity). In similar panels in subsequent figures, intermediate colors indicate relative tendencies toward each phenotype. (D) Seam cell polarity phenotypes of compound receptor mutants. All strains in this and subsequent figures carry *elt-3p*::GFP. *mom-5(AID)* refers to *mom-5(os171) osTi8*[*eft-3p*::TIR1::mRuby] animals grown on plates containing 250 µM IAA. § Data from (Yamamoto et al. 2011). † Homozygotes derived from heterozygous mothers. The sum of percentages may not equal 100 because values were rounded to the nearest whole number, and divisions along the dorsoventral axis (indicated by asterisks) were excluded from percentage calculations.

Using this strategy, we previously showed that seam cell (V1‒V5) polarity is randomized in *3xWnt (cwn-1 egl-20 cwn-2)* mutants (Yamamoto et al. 2011). Notably, the polarity distributions obtained from *elt-3p*::GFP‒based scoring closely paralleled those determined by GFP::POP-1/TCF nuclear localization. In addition, mutants defective in canonical Wnt signaling (*pry-1*/Axin and *bar-*1/β-catenin) did not show defects in asymmetric seam daughter fates (n = 20 at the L1 stage for each genotype). Therefore, in Wnt and receptor compound mutants described below, seam cell phenotypes are unlikely to be influenced by direct regulation of cell fates through the canonical pathway, but instead primarily reflect defects in Wnt-dependent cell polarity. Accordingly, *elt-3p*::GFP provides a reliable readout of seam cell polarity.

In our previous study, we also showed that three receptors̶LIN-17/Fzd, MOM-5/Fzd, and CAM-1/Ror̶redundantly regulate seam cell polarity (Yamamoto et al. 2011). Single or double mutants of these receptors exhibited minor or moderate polarity defects, which were strongly enhanced in triple receptor mutants (Figure 1D). Because *mom-5 cam-1* double null mutants are zygotically embryonic lethal, we previously analyzed *mom-5(null)* homozygotes that retained potential maternal contribution from heterozygote mothers and reduced *cam-1* by RNAi, which might have caused only partial knockdown. To obtain clearer results, we inhibited *mom-5* using the auxin-inducible degron (AID) system (Nishimura et al. 2009; Zhang et al. 2015) in the *cam-1(gm122)* null background. We inserted AID::mClover at the C terminus of the endogenous *mom-5* locus to generate *mom-5(os171)*.

When *mom-5(os171) cam-1(gm122)* animals were grown on plates containing DMSO from late embryonic stages, only minor polarity defects were observed (Supplementary Figure 1). In contrast, when grown on plates containing auxin (IAA) dissolved in DMSO, they exhibited much stronger defects than *mom-5(null) cam-1(RNAi)* animals (Figure 1D). Hereafter, animals carrying *mom-5(os171)* grown on IAA plates are referred to as *mom-5(AID)*. In *mom-5(AID) cam-1(gm122)* animals, polarity of the anterior seam cells (V1‒V4) was almost completely lost, whereas that of the posterior V5 and V6 cells remained nearly normal. Addition of a *lin-17(n3091 null)* mutation caused symmetric divisions of all (V1‒V6) seam cells (*lin-17 mom-5(AID) cam-1*). These results suggest that anterior and posterior seam cells may rely on distinct receptor combinations to establish polarity. However, because the polarity defects of V1‒V4 in *mom-5(null) cam-1(RNAi)* were strongly enhanced by the *lin-17* mutation (Figure 1D) (Yamamoto et al. 2011), *lin-17* can regulate V1‒V4 polarity when *cam-1* function is compromised but not completely lost. The weaker requirement of LIN-17 in anterior cells may reflect its lower expression there (Yamamoto et al. 2011; Ji et al. 2013). Consistent with this, expressing LIN-17 in all seam cells using the seam-cell specific *scm* promoter rescued polarity loss in *mom-5(AID) cam-1(gm122)*. However, polarity orientation became random, resembling *3×Wnt* mutants. Thus, LIN-17 alone can polarize seam cells, whereas proper orientation of anterior cells requires additional receptors. In contrast, the normal polarity of V5 and V6 in *mom-5(AID) cam-1* indicates that polarity orientation of these posterior cells is controlled solely by LIN-17. Since *lin-17(null) cam-1(null)* with normal MOM-5 function show nearly normal polarity of seam cells except for V6 (Yamamoto et al. 2011), seam cells are polarized when either of the Frizzled receptors, LIN-17 or MOM-5, is present at a sufficient level.

The use of *mom-5(AID)* enabled us to examine the function of the C-terminal kinase domain of CAM-1 using the *cam-1(ks52)* allele, which produces a receptor lacking this domain (Koga et al. 1999). Although CAM-1 protein level is reduced in the *ks52* mutant (Chien et al. 2015), *cam-1(ks52)* shows much less severe effects than *cam-1(gm122 null)* in multiple neuronal phenotypes (Kim and Forrester 2003; Francis et al. 2005; Kennerdell et al. 2009), indicating that *ks52* is a hypomorphic mutation. We found that *mom-5(AID) cam-1(ks52)* and *lin-17 mom-5(AID) cam-1(ks52)* showed similar defects to *mom-5(AID) cam-1(gm122 null)* and *lin-17 mom-5(AID) cam-1(gm122*). These results suggest that the C-terminal kinase domain plays an important role for CAM-1 function in polarity regulation.

We found that *mom-5(AID)* caused stronger defects than *mom-5(ne12 null)* from heterozygote mothers in the *lin-17(n3091 null)* background. For example, the loss of polarity of V6 was 100% in *lin-17 mom-5(AID)* animals, compared to 17% in *lin-17(null) mom-5(null)* from heterozygotes, indicating strong maternal effects of *mom-5*, which is expressed in and essential for early embryogenesis (Rocheleau et al. 1997; Thorpe et al. 1997; Heppert et al. 2018). This is surprising, since we could not detect fluorescence from maternally derived *mom-5*::mNeonGreen (mNG) *(cp367)* (Heppert et al. 2018) in *+/+* L1 larvae from heterozygote mothers (*+/cp367*) (data not shown). Therefore, undetectable amounts of maternal MOM-5 can regulate polarity.

### Genetic interactions between Wnts and receptors

To analyze the relationship between Wnts and receptors, we combined single or double receptor mutations with single, double, or triple Wnt mutations and analyzed seam cell polarity (Figures 2 and 3). We describe the results focusing on the function of each receptor. We have shown that three Wnts (CWN-1, EGL-20, and CWN-2) redundantly control seam cell polarity, since any single or double Wnt mutants show only minor polarity defects except for V5, which is affected in *egl-20* single mutants (Yamamoto et al. 2011) (Figure 2, first column). Although V6 polarity is nearly normal in *3xWnt*, it is redundantly regulated by four Wnt genes (*lin-44*, *cwn-1*, *egl-20*, and *cwn-2*) and disrupted in the quadruple Wnt mutants. Therefore, if single or double Wnt mutations cause strong polarity defects in the absence of one of the receptors, the non-mutated Wnt’s function is likely to depend on the mutated receptor.

**Figure 2.**
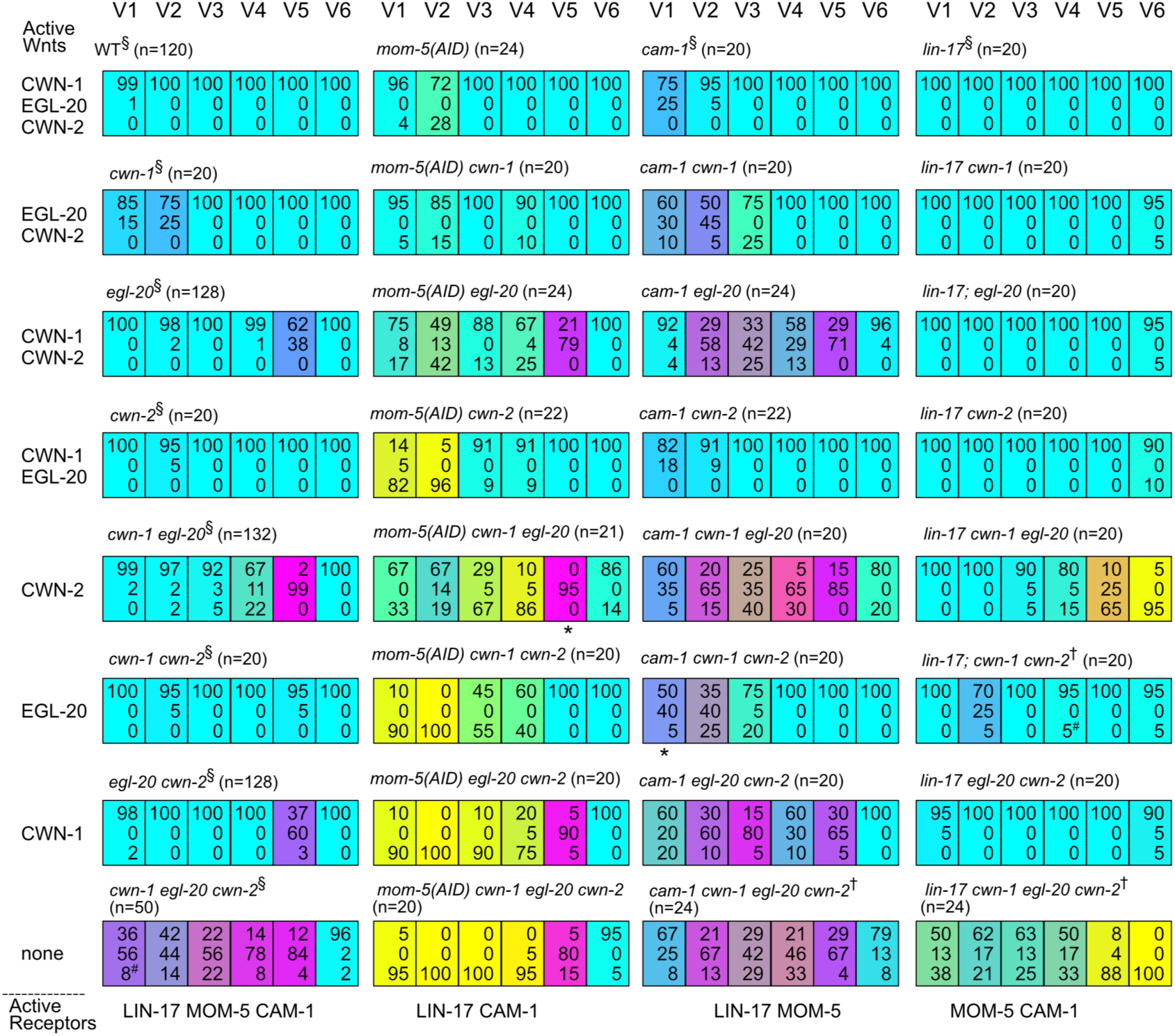
Analyses of compound mutants containing a single receptor mutation. Each colored box represents the polarity of individual seam cell divisions, as described in Figure 1C. On the left side of each column and below each row, the active Wnts and receptors of the compound mutants, respectively, are shown. § Data from (Yamamoto et al. 2011). †Homozygotes derived from heterozygous mothers. # Includes one symmetric division that produced two seam daughters. The sum of percentages may not equal 100 because values were rounded to the nearest whole number, and divisions along the dorsoventral axis (indicated by asterisks) were excluded from percentage calculations.

**Figure 3.**
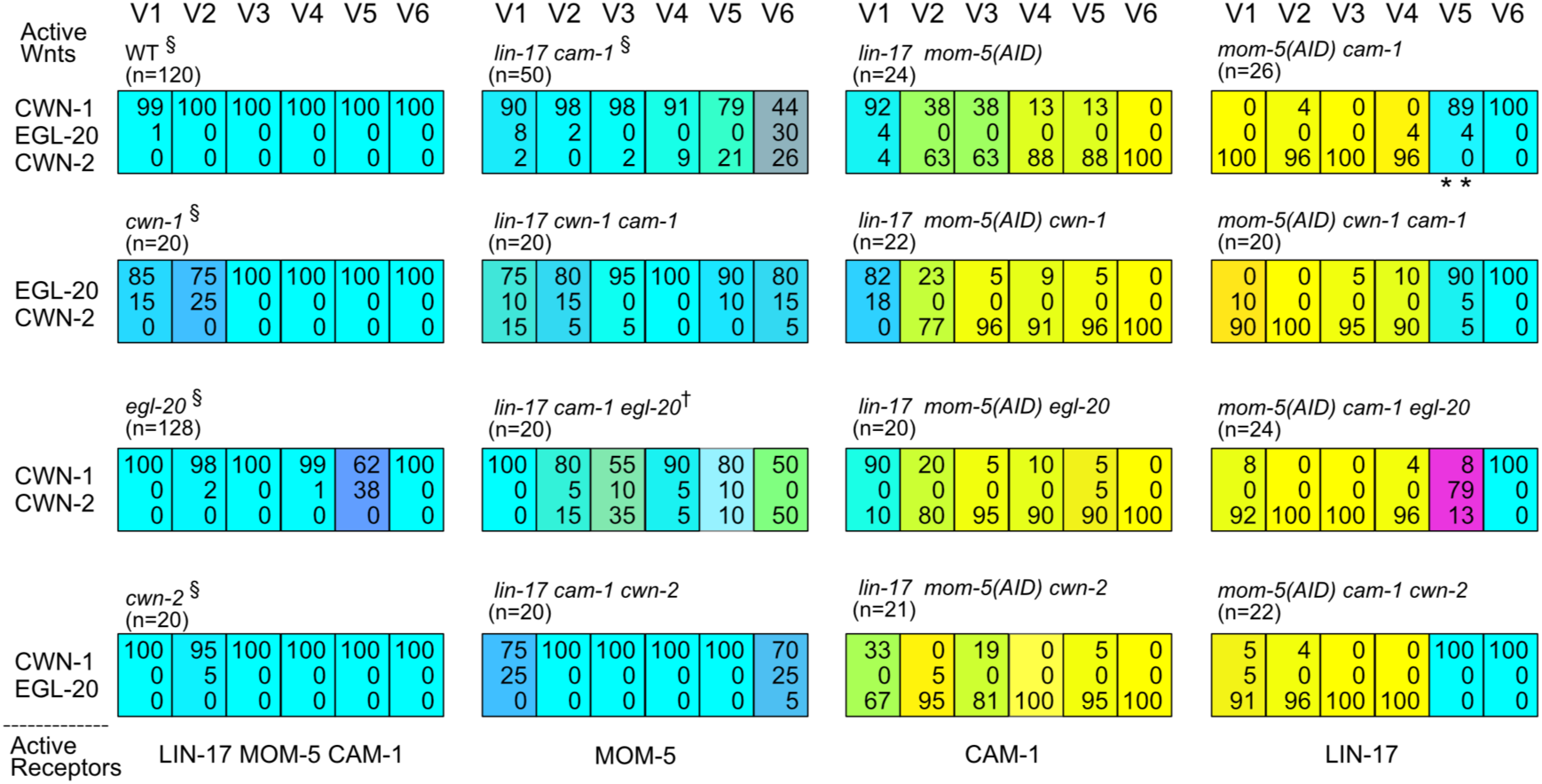
Analyses of compound mutants containing double receptor mutations. Each colored box represents the polarity of individual seam cell divisions, as described in Figure 1C. On the left side of each column and below each row, the active Wnts and receptors of the compound mutants, respectively, are shown. §Data from (Yamamoto et al. 2011). †Homozygotes derived from heterozygous mothers. The sum of percentages may not equal 100 because values were rounded to the nearest whole number, and divisions along the dorsoventral axis (indicated by asterisks) were excluded from percentage calculations.

### MOM-5 regulates cell polarization in response to all three Wnts

In the *mom-5(AID)* background, *cwn-2* mutations strongly disrupted polarity in V1/V2 (p < 0.01 in all cases compared to *cwn-2*) (Figure 2, second column), suggesting that both CWN-1 and EGL-20 functions are compromised in the absence of *mom-5*. Similarly, moderate polarity defects in V2/V4 (p < 0.01 in all cases compared to *egl-20*) in *mom-5(AID) egl-20* suggest that the functions of CWN-1 and CWN-2 are also affected in *mom-5(AID)*. Consistently, moderate to strong polarity defects in any double Wnt mutants in the *mom-5(AID)* background indicate that all Wnt functions depend, at least in part, on *mom-5*. Since *mom-5(AID) egl-20 cwn-2* showed the strongest defects among them, CWN-1 appears to depend more strongly on MOM-5 than EGL-20 or CWN-2. However, because the *cwn-1* mutation enhanced the defects of V3/V4 in *mom-5(AID) egl-20* and *mom-5(AID) cwn-2* (p < 0.05 in all cases), CWN-1 has MOM-5 independent function as well. In the *lin-17 cam-1* background, in which MOM-5 is the only remaining receptor, single Wnt mutations did not cause strong phenotypes (Figure 3, second column), confirming redundant functions of the three Wnts through MOM-5. In summary, MOM-5 mediates the functions of all three Wnt genes. Because the major defects in any compound mutants containing *mom-5(AID)* are loss of polarity, MOM-5 plays a central role in cell polarization, although it also regulates polarity orientation, considering nearly normal polarity orientation in *lin-17 cam-1* animals.

### CWN-2 functions mostly through CAM-1, which regulates polarity orientation

The *cam-1 cwn-1 egl-20* triple mutants showed strong phenotype similar to *cam-1 3xWnt*, suggesting that CWN-2 function strongly depends on CAM-1. Consistently, the *cwn-2* mutation had only minor effects in the *cam-1*, *cam-1 cwn-1*, *cam-1 egl-20* backgrounds (Figure 2, third column). Although the *cwn-1* mutation caused significant enhancements in V2 (p < 0.01) and V3 (p < 0.05) in *cam-1*, indicating that CWN-1 retains some activity to control polarity in the absence of *cam-1*, the similar defects between *cam-1 3xWnt*s and *cam-1 egl-20 cwn-2* suggest that CWN-1 function is largely disrupted in *cam-1*. In contrast, strong enhancements caused by the *egl-20* mutation in *cam-1* (V2‒V5), *cam-1 cwn-1* (V3‒V5), *cam-1 cwn-2* (V2‒V5), and *cam-1 cwn-1 cwn-2* (V2‒V5) (p < 0.01 in all cases) indicate that EGL-20 retains strong activity in *cam-1*. Since *egl-20* alone was not sufficient for normal polarity in V1‒V3 of *cam-1 cwn-1 cwn-2* (p < 0.05 in all cases compared to *cwn-1 cwn-2*), EGL-20 function partly depends on *cam-1* in these cells, which are located far from the EGL-20 source.

In the *lin-17 mom-5(AID)* background, in which CAM-1 is the only remaining major receptor, the *cwn-2* mutation caused stronger polarity defects than *egl-20* or *cwn-1* mutations (p < 0.01 in V1) (Figure 3, third column). Both *cwn-1* and *egl-20* moderately enhanced the *lin-17 mom-5(AID)* phenotype in V2 (p < 0.05), suggesting that CAM-1 can act as a receptor for these Wnts, at least in the absence of LIN-17 and MOM-5 functions. Taken together, these results indicate that although CAM-1 is capable of mediating signaling from all three Wnts, the degree of dependency varies among them̶strongest for CWN-2, moderate for CWN-1, and weakest for EGL-20.

In contrast to the *mom-5* backgrounds, in which loss of polarity was more frequent than reversal (Figure 2, second column), the *cam-1* background primarily showed polarity reversal as the major defect (Figure 2, third column). For example, *cam-1 egl-20 cwn-2* exhibited random polarity in the V1‒V5 cells, while *cam-1 cwn-1 cwn-2* showed random polarity in V1 and V2. Thus, in the absence of *cam-1*, *cwn-1* and *egl-20* cannot properly regulate the orientation of these cells, indicating that CAM-1 plays a crucial role in controlling polarity orientation.

### LIN-17 may function as a switch for polarity orientation

The *lin-17* mutation caused strong defects in V6 in the *cwn-1 egl-20* double mutant (p < 0.01) (note that V5 is already abnormal in *cwn-1 egl-20*), suggesting that CWN-2 function depends on LIN-17, at least in V6, which is located far from the CWN-2 source cells (Figure 2, fourth column). The strong V6 phenotype in *lin-17 cwn-1 egl-20* further suggests that LIN-44/Wnt, expressed in the tail hypodermal cells and known to act redundantly with these Wnts in V6 (Yamamoto et al. 2011), also depends on LIN-17, as previously shown for the T cell (a seam cell posterior to V6) (Herman et al. 1995; Goldstein et al. 2006).

In the *mom-5(AID) cam-1* background, where LIN-17 is the only remaining receptor among the three, the *egl-20* mutation caused strong polarity reversal in V5 (p < 0.01), suggesting that EGL-20 controls V5 polarity orientation through LIN-17 (Figure 3, fourth column). In contrast to EGL-20 and CWN-2, which both act at least partly through LIN-17, CWN-1 function appears largely independent of LIN-17, as indicated by the nearly normal polarity in *lin-17 egl-20 cwn-2*, where only CWN-1 is expressed. Moreover, the *cwn-1* mutation had no effect on V5/V6 polarity in the *mom-5(AID) cam-1* background, further suggesting that LIN-17 alone cannot mediate CWN-1 function.

Polarity reversal of V5 caused by the *egl-20* single mutation was suppressed by the *lin-17* mutation (p<0.01), as reported previously (Whangbo et al. 2000). Similarly, the *lin-17* mutation suppressed polarity reversal of V5 in most compound mutants showing strong reversal phenotypes (V5 in *cwn-1 egl-20*, V2‒V3 and V5 in *cam-1 egl-20*, and V1‒V5 in the *3xWnt* mutants; p<0.05 in all cases) (Figure 2 and 3), indicating that *lin-17* promotes reversed polarity in these genetic backgrounds. In contrast, in the absence of the *egl-20* mutation, *lin-17* promotes normal polarity in V1-V6 in *mom-5(null) cam-1(RNAi)* (p<0.01 in all cases) and in V2‒V6 in *mom-5(AID)* (p<0.05 in V2 and p<0.01 in V3‒V6) (Figure 1). Therefore, LIN-17 may function as a molecular switch for polarity orientation, promoting either normal or reversed polarity depending on whether EGL-20/Wnt is present (and possibly activates the LIN-17 receptor) or absent.

### Wnt-independent polarization by LIN-17 and MOM-5

We have shown that seam cells are mostly polarized in random orientations in *3xWnt* mutants as well as in quintuple Wnt mutants carrying mutations in all Wnt genes (Yamamoto et al. 2011) (Figure 2, last row), suggesting Wnt-independent polarization. We found that *lin-17 3xWnt* exhibited strong loss of polarity in V5 and V6. Complementarily, *mom-5 3xWnt* showed loss of polarity in V1‒V4. These results indicate that Frizzled receptors (either MOM-5 or LIN-17) are essential for Wnt-independent polarization. The position-dependent effects may reflect posteriorly biased expression of LIN-17 (Yamamoto et al. 2011; Ji et al. 2013). In contrast to MOM-5 and LIN-17, CAM-1 appears to play only minor roles in Wnt-independent polarity, as *cam-1 3xWnt* displayed defects similar to those of *3xWnt*. Our findings suggest that Frizzled receptors possess Wnt-independent activities that polarize cells, whereas CAM-1/Ror primarily mediates Wnt-dependent polarization.

### EGL-20 and CWN-1 have both instructive and permissive functions

We previously showed that reversing the gradients of *cwn-1* and *cwn-2* suppresses the polarity defects of *3xWnt* mutants, indicating that Wnts can act permissively (Yamamoto et al. 2011). In those experiments, the *cwn-2* gradient was reversed by posterior expression of CWN-2::Venus (a C-terminal fusion) under the *egl-20* promoter. However, we later found that a C-terminal mNG fusion of endogenous *cwn-2* disrupts its function (see Materials and Methods). Therefore, to reverse the *cwn-2* gradient in the present study, we used an N-terminal fusion (*egl-20p*::mNG::CWN-2). This construct strongly suppressed the polarity defects of *3xWnt* in V1‒V4 (p < 0.01 for all cells), confirming the permissive activity of *cwn-2* (Figure 4). Similarly, reversing the *egl-20* gradient using *ceh-22p*::EGL-20::mNG (*ceh-22* is expressed in the pharynx) suppressed the *3xWnt* phenotype in V2‒V4 (p < 0.05 for all cells), supporting permissive functions of both Wnts.

**Figure 4.**
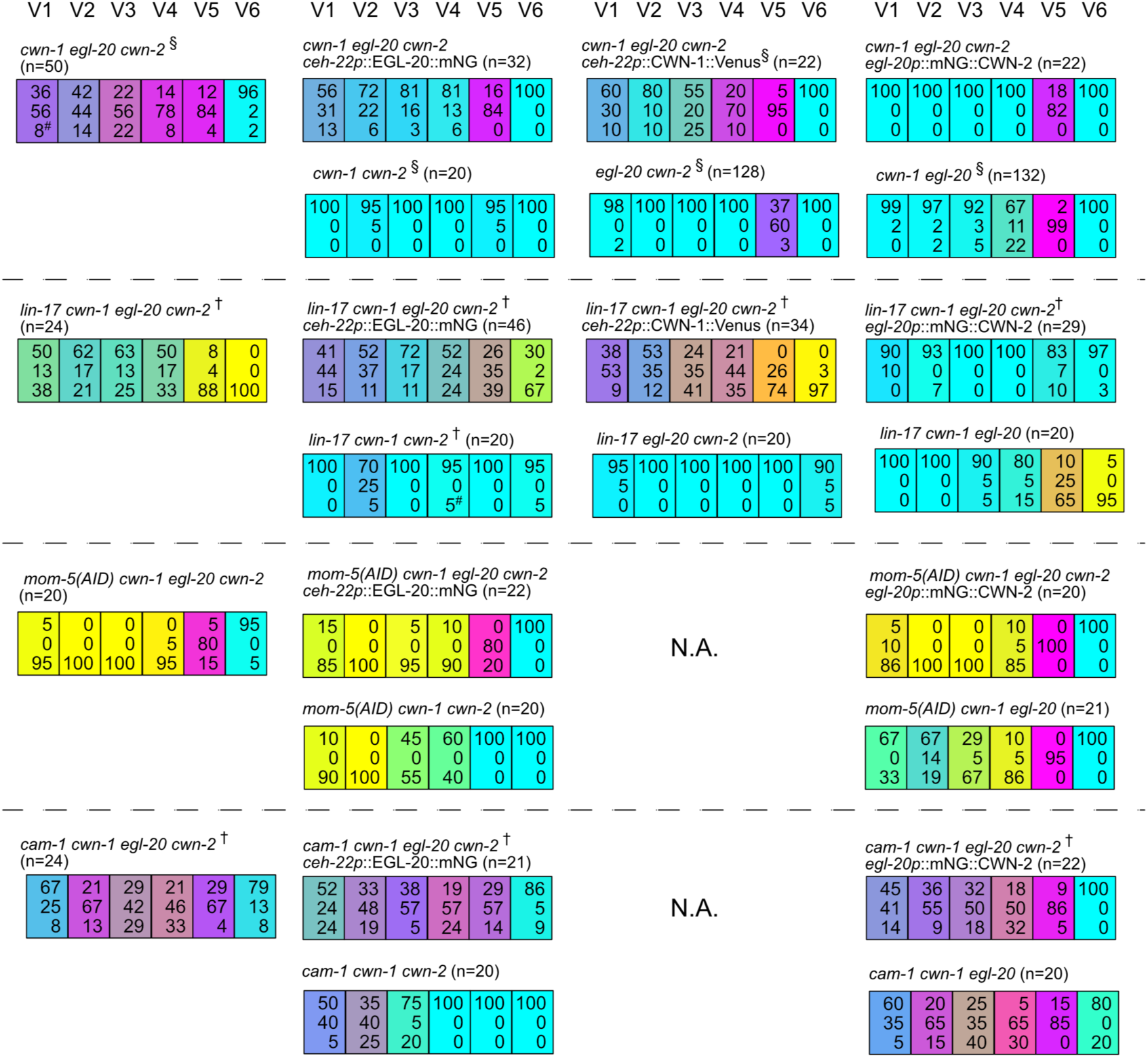
CWN-1 and EGL-20 have both instructive and permissive functions. Each colored box represents the polarity of individual seam cell divisions as described in Figure 1C. When the ectopic expression of a particular Wnt fully compensated for the loss of that Wnt in the corresponding compound mutant, the resulting phenotype is expected to match that shown below each ectopic expression dataset. N.A., not analyzed because of technical limitations. § Data from (Yamamoto et al. 2011). #Includes one symmetric division that produced two seam daughters. † Homozygotes derived from heterozygous mothers. The sum of percentages may not equal 100 because values were rounded to the nearest whole number.

We next analyzed the effects of reversing Wnt gradients in strains carrying receptor mutations in addition to the *3xWnt* mutations. Contrary to the rescue of polarity defects observed in *3xWnt* mutants, we unexpectedly found that *ceh-22p*::CWN-1::Venus significantly enhanced polarity reversal in *lin-17 3xWnt* (p < 0.01 in V1; p < 0.05 in V4 and V5; p > 0.05 in V2 and V3, but p < 0.05 when V2 and V3 were combined). Similarly, *ceh-22p*::EGL-20::mNG enhanced polarity reversal in *lin-17 3xWnt* at least in V1 (p < 0.05) and V5 (p < 0.01). These results suggest that the *egl-20* and *cwn-1* gradients instructively control polarity orientation, at least in the absence of LIN-17. In contrast, *egl-20p*::mNG::CWN-2 strongly rescued polarity defects in *lin-17 3xWnt* as it did in *3xWnt* strains (p < 0.01 in all cases).

In contrast to the enhancement of polarity reversal observed in the *lin-17 3xWnt* background, *ceh-22p*::EGL-20::mNG showed only minor effects in the *mom-5(AID) 3xWnt* and *cam-1 3xWnt* backgrounds. Because *ceh-22p*::EGL-20::mNG exhibits only weak activity even in the *3xWnt* background, these results suggest that both *mom-5* and *cam-1* are required to facilitate the limited function of EGL-20 expressed under the *ceh-22* promoter. Similarly, *egl-20p*::mNG::CWN-2 produced only minor effects in the *mom-5(AID) 3xWnt* and *cam-1 3xWnt* backgrounds. The weak effects in the *cam-1* background are consistent with the results described above that CWN-2 function strongly depends on *cam-1*. However, given the strong rescuing activity of *egl-20p*::mNG::CWN-2 in the *3xWnt* background, its weak effects in the *mom-5(AID) 3xWnt* background were unexpected. Since *mom-5(AID) cwn-1 egl-20* mutants, in which endogenous anterior CWN-2 is active, showed milder defects in V1‒V3, posteriorly expressed CWN-2 may be less effective than the endogenous source, possibly due to the greater distance between V1‒V3 and the CWN-2‒expressing cells.

### DSH proteins are required for seam cell polarity

DSH/Dishevelled proteins are conserved intracellular signaling components that act downstream of Frizzled receptors. *C. elegans* has three homologs: DSH-1, DSH-2, and MIG-5. We previously showed that two DSH proteins, DSH-2 and MIG-5, localize asymmetrically during seam cell divisions, as in planar cell polarity (PCP) regulation in other organisms (Mizumoto and Sawa 2007a). However, their functional requirements in seam cell polarity have not been demonstrated. We found that double deletion mutants, *dsh-2 dsh-1* and *dsh-2 mig-5*, exhibited nearly normal seam cell polarity (Figure 5), suggesting redundant functions among the three homologs. Triple deletion mutants were nearly embryonic lethal (data not shown). Therefore, we utilized the previously reported mNG::AID-tagged *mig-5* allele *(cp385)* (Heppert et al. 2018) to generate *dsh-2 dsh-1 mig-5(AID)* triple mutants. Growing these mutants on auxin-containing plates caused a strong loss of polarity (p < 0.01 in V1-V6), indicating that DSH proteins are required for cell polarization, consistent with their conserved role in PCP regulation across species.

**Figure 5.**
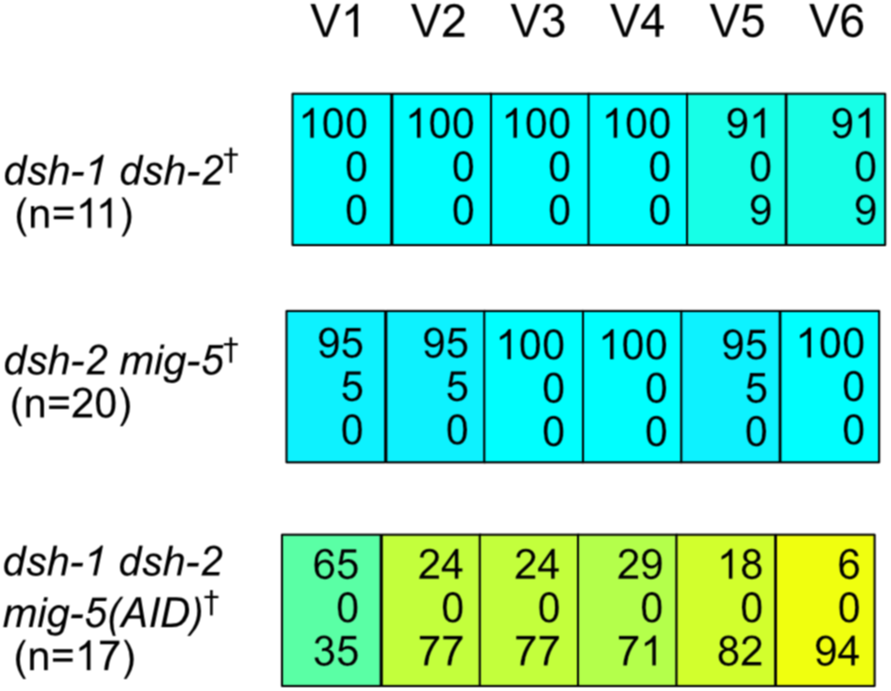
DSH proteins are required for seam cell polarity. Each colored box represents the polarity of individual seam cell divisions as in Figure 1C. *mig-5(AID)* refers to animals containing *mig-5(cp385) osTi9*[*eft-3p*::TIR1::mRuby] grown on plates containing 250 µM IAA. † Homozygotes derived from heterozygous mothers. The sum of percentages may not equal 100 because values were rounded to the nearest whole number.

## Discussion

### Non-redundant functions of Wnt receptors

Our results demonstrate that, although three Wnt ligands and three receptors redundantly regulate seam cell polarity, specific Wnt‒receptor preferences are evident. The function of CWN-2 strongly depends on CAM-1, as the *cwn-2* mutation had only minor effects in strains carrying a *cam-1* mutation. However, CWN-2 function was also compromised by *mom-5* or *lin-17* mutations in certain genetic backgrounds, such as the V2/V4 cells in *mom-5(AID) egl-20* and the V6 cell in *lin-17 cwn-1 egl-20*. One possibility is that CAM-1/Ror acts as a co-receptor with Frizzleds (LIN-17 or MOM-5). Although this co-receptor model may seem inconsistent with the effects of *cwn-2* mutation in *lin-17 mom-5(AID)*, particularly in the V1 cell, other Frizzled receptors such as MIG-1/Fzd and CFZ-2/Fzd may compensate for the loss of LIN-17 and MOM-5.

Consistent with our results, roles of CWN-2 and CAM-1 have also been reported in the nervous system of *C. elegans*. CWN-2 regulates the position of the nerve ring through CAM-1 and the Frizzled receptors CFZ-1 and MIG-1 (Kennerdell et al. 2009). It also controls neurite outgrowth of the RME neuron via CAM-1 (Song et al. 2010). In addition, CWN-2 expressed in cholinergic motor neurons regulates acetylcholine receptor translocation and synaptic plasticity through the CAM-1/LIN-17 receptor complex in adult animals (Jensen et al. 2012). Therefore, CAM-1 is likely to function as a main receptor for CWN-2 in both nervous and epithelial cells.

In contrast to CWN-2, the functions of CWN-1 and EGL-20 depend strongly on MOM-5 and partially on CAM-1. In the *mom-5(AID) cam-1* background, EGL-20̶but not CWN-1̶plays an essential role in V5 polarity, indicating that EGL-20, but not CWN-1, activates LIN-17 to regulate proper polarity orientation in the absence of MOM-5 and CAM-1.

### Regulation of polarity orientation

Among the polarity phenotypes observed in the various compound mutants we analyzed, random polarity orientation occurred only in *cam-1* mutant backgrounds, except for the *3xWnt* mutant, suggesting that CAM-1 plays a crucial role in regulating polarity orientation. In addition, LIN-17 appears to be a key component in this regulation. The suppression of polarity reversal in strains carrying the *egl-20* mutation by the *lin-17* mutation suggests that LIN-17 determines polarity orientation depending on activation by EGL-20. Although this model seems inconsistent with the finding that uniform LIN-17 expression by the *scm* promoter in *mom-5(AID) cam-1* caused random polarity orientation in V1‒V4 cells even in the presence of active EGL-20, this apparent contradiction can be explained if regulation of polarity orientation by LIN-17 requires either MOM-5 or CAM-1. In their absence, LIN-17 may promote random polarity. In addition to LIN-17 and CAM-1, MOM-5 also contributes to polarity orientation as well as polarization, since seam cell polarity was nearly normal in *lin-17 cam-1* double mutants.

### A possible model for the regulation of polarity orientation

Although the mechanism of polarity regulation remains unclear̶particularly because polarity cues for permissive Wnt functions are not yet known̶our genetic analyses of Wnts and their receptors allow us to propose possible models for their functions (Figure 6). We have shown that Wnts possess both instructive and permissive roles in regulating polarity orientation. Considering that instructive effects were observed only in the *lin-17* background, *lin-17* may either inhibit the instructive effects or enhance the permissive effects. The former possibility seems inconsistent with the observation that *lin-17* promotes normal polarity orientation in the presence of Wnts. Therefore, it is more likely that the permissive effects of Wnts are at least partially impaired in *lin-17* mutants.

**Figure 6.**
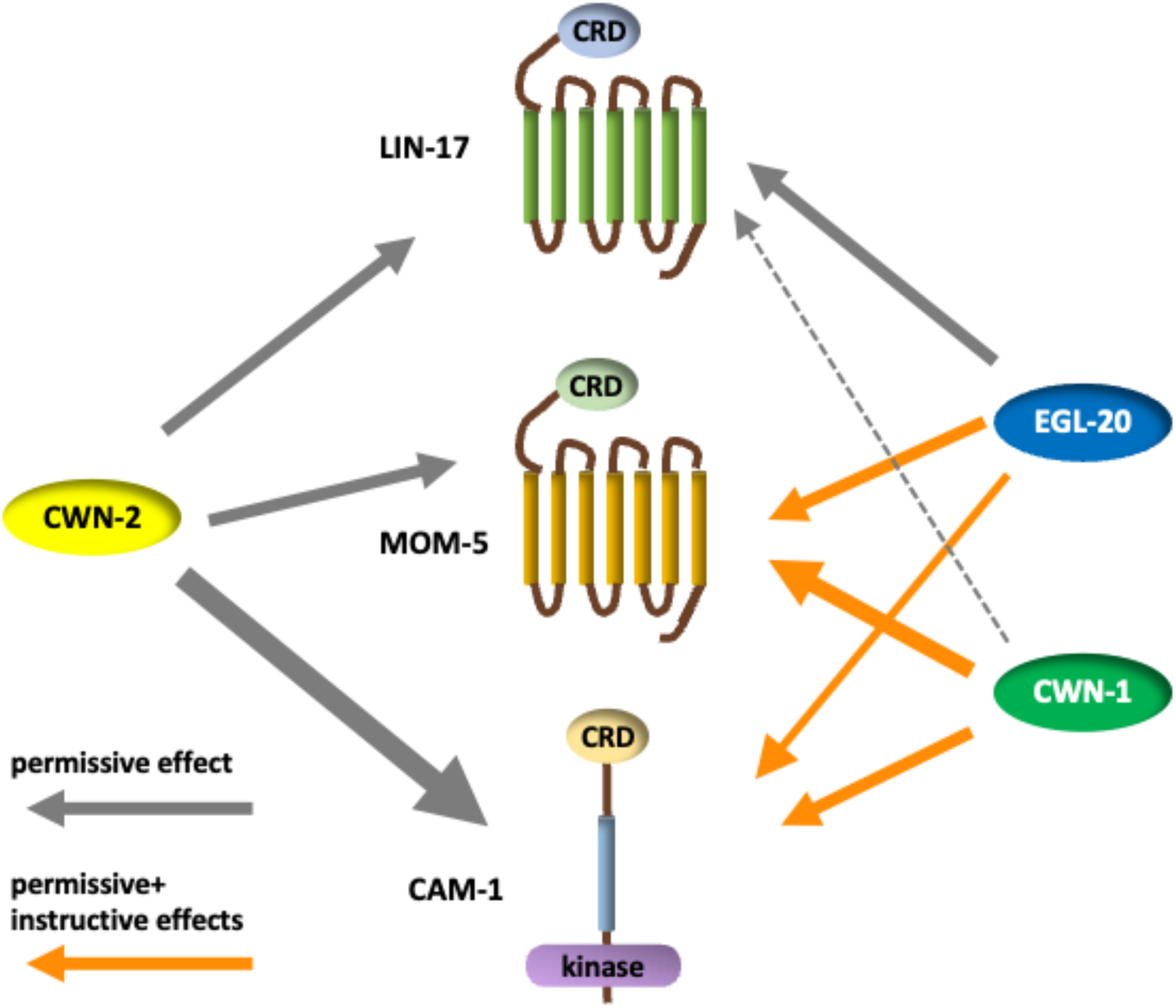
A possible model for polarity regulation by Wnts and their receptors. Gray and orange arrows indicate permissive and combined permissive + instructive effects of Wnt proteins, respectively. Arrow thickness represents the relative dependence of each Wnt function on its receptors.

For the permissive effects, polarity orientation is determined by Wnt-gradient‒independent cues. The random polarity orientation induced by uniform LIN-17 expression in the *mom-5(AID) cam-1* background indicates that *lin-17* itself̶or its activation by Wnts̶ does not provide such cues. Instead, *mom-5* and/or *cam-1*, which also contribute to polarity orientation, may mediate these cues and act together with *lin-17* to orient cell polarity correctly.

The Wnt-gradient‒independent cues require Wnt activity to regulate polarity orientation. Although all three Wnts can function permissively, *lin-17* may be required for the permissive functions of *cwn-1* and *egl-20*, but not for *cwn-2*, since ectopic expression of *cwn-2* rescued the *lin-17 3xWnt* phenotype. In contrast, the permissive effects of ectopic expression of all Wnts depend on *cam-1* and *mom-5*. Therefore, *cwn-2* may exert its permissive effect through *mom-5* and/or *cam-1*. Given its strong dependence on *cam-1*, *cam-1* is the most plausible target of *cwn-2*. Similarly, the permissive effects of *cwn-1* and *egl-20* may act through *mom-5* and/or *cam-1*, in addition to *lin-17* (in the case of *egl-20*), consistent with the nearly normal polarity observed in *lin-17* single and *lin-17 cam-1* double mutants.

The instructive effects of *cwn-1* and *egl-20* are likely mediated by *mom-5* and *cam-1*, since they were observed in the *lin-17* background. The absence of instructive effects of *cwn-2* is reasonable, given that its gradient is oriented opposite to those of *cwn-1* and *egl-20*. The strong requirement for *cam-1* may be key to understanding the distinct function of *cwn-2* compared with *cwn-1* and *egl-20*. To elucidate how polarity orientation is determined, it will be necessary to identify the polarity cues that underlie the permissive Wnt functions. In other organisms, the roles of Wnts in polarity regulation have been analyzed under the assumption that they are either instructive or permissive. However, Wnts may in fact possess both instructive and permissive functions in polarity regulation, as demonstrated here in *C. elegans*.

## Data availability

All strains are listed in Supplementary Table S1 and available upon request. The author affirms that all data necessary for confirming the conclusions of the article are present within the article, figures, and tables.

## Acknowledgements

We thank Masato Kanemaki for the advices on the auxin degron experiments, Jennifer Heppert and Bob Goldstein for plasmids used for *mom-5* genome editing, Mayumi Onami for technical helps and generating a transgenic allele used in the study, Tazhibayeva Samal for comments on the manuscript. Some strains were provided by the CGC, which is funded by NIH Office of Research Infrastructure Programs (P40 OD010440), and by the National Bioresource Project of Japan (NBRP).

---

**Table S1.**
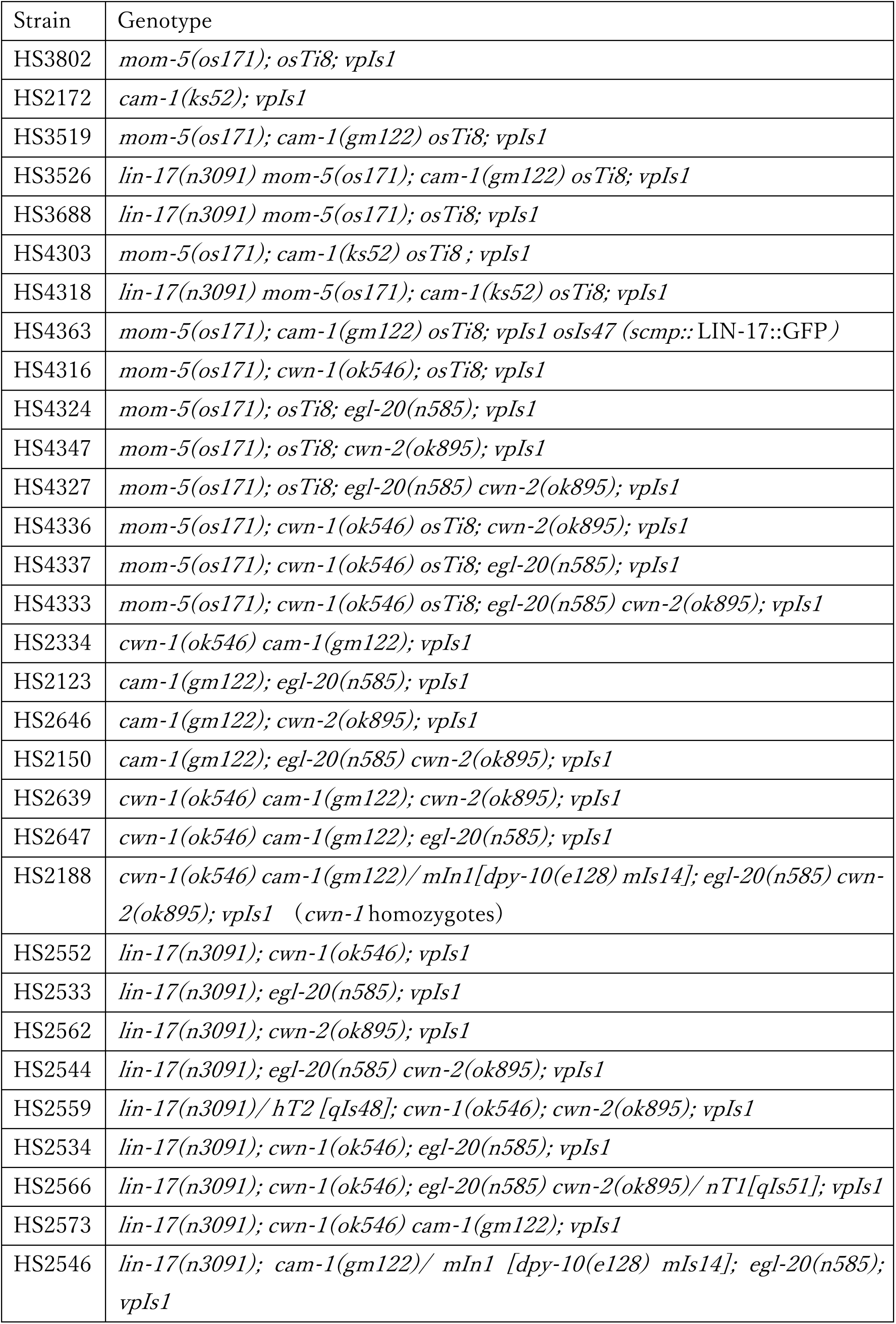
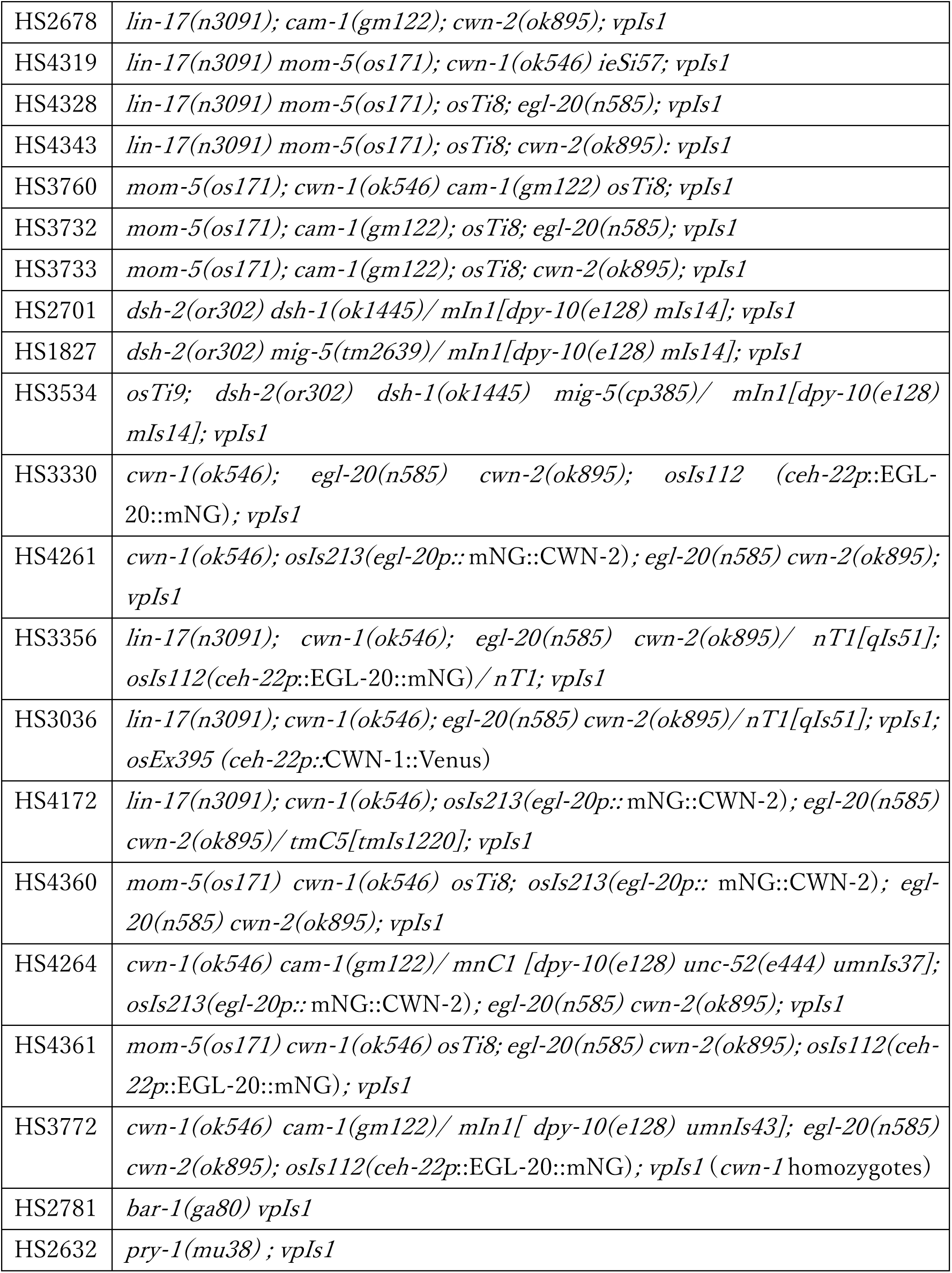
List of strains used for experiments.

**Supplemental Figure 1.**
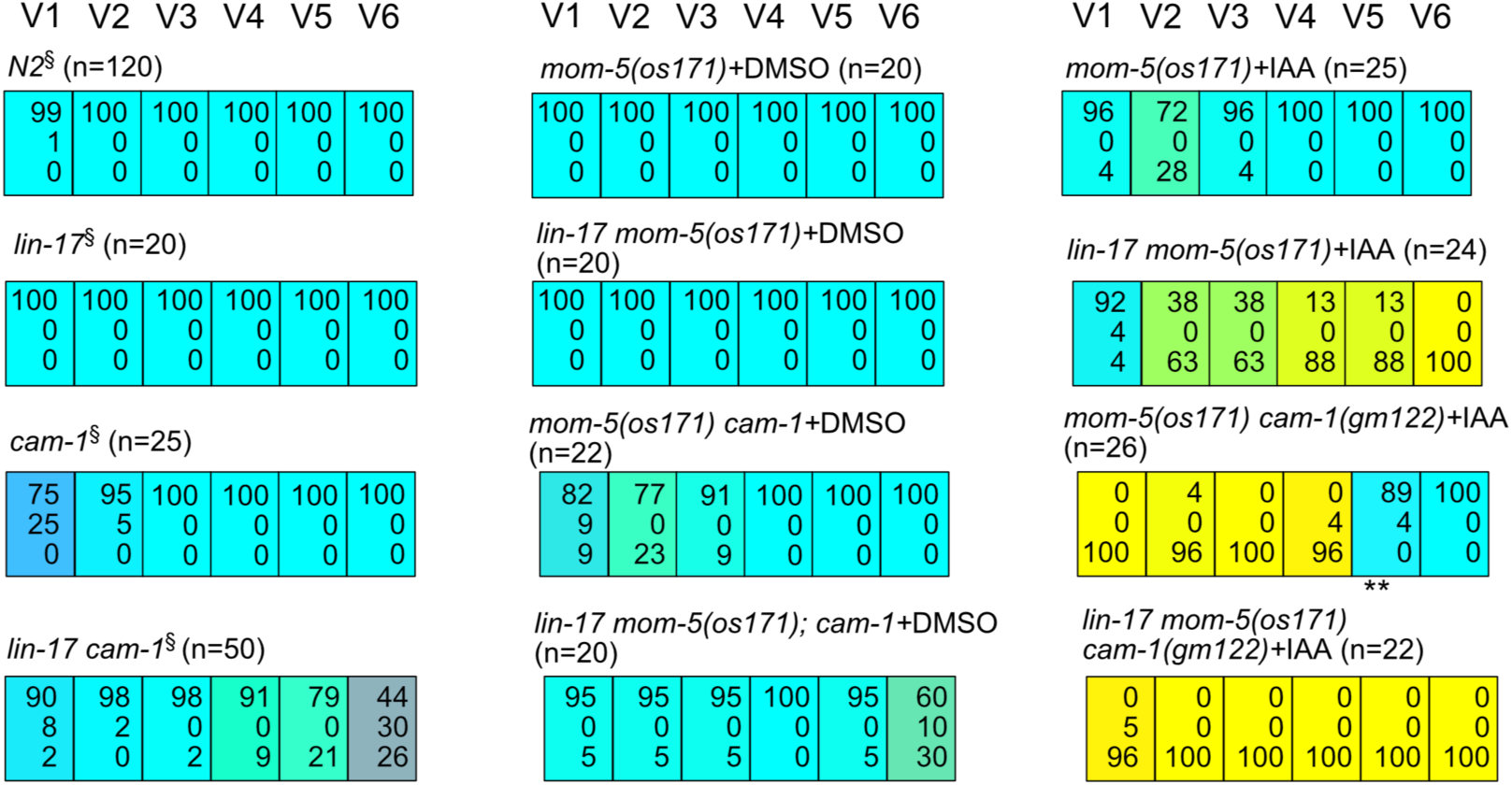
Specificity of auxin treatments on *mom-5(AID)* Each colored box represents the polarity of individual seam cell divisions as in Figure 1C. Animals containing *mom-5(os171) osTi8*[*eft-3p*::TIR1::mRuby] were grown on plates containing either 0.05% DMSO or 250μM IAA dissolved in DMSO (0.05%). §Data from (Yamamoto et al. 2011). The sum of percentages may not equal 100 because values were rounded to the nearest whole number, and divisions along the dorsoventral axis (indicated by asterisks) were excluded from percentage calculations.

